# Nociception in chicken embryos, Part III: Analysis of movements before and after application of a noxious stimulus

**DOI:** 10.1101/2023.04.20.537674

**Authors:** Stephanie C. Süß, Julia Werner, Anna M. Saller, Larissa Weiss, Judith Reiser, Janie M. Ondracek, Yury Zablotski, Sandra Kollmansperger, Malte Anders, Benjamin Schusser, Thomas Fenzl, Christine Baumgartner

**Affiliations:** Center for Preclinical Research, TUM School of Medicine, Technical University of Munich, Germany; Chair of Zoology, TUM School of Life Sciences Weihenstephan, Technical University of Munich; Clinic for Swine, Center for Clinical Veterinary Medicine, LMU Munich, Bavaria, Germany; Clinic for Anesthesiology and Intensive Care, TUM School of Medicine, Technical University of Munich, Germany; Reproductive Biotechnology, TUM School of Life Sciences, Technical University of Munich, Weihenstephan, Technical University of Munich, Germany

## Abstract

Many potentially noxious interventions are performed on chicken embryos in research and in the poultry industry. It is therefore in the interest of animal welfare to define the point at which a chicken embryo is capable of nociception. The present part III of a comprehensive study examined the movements of developing chicken embryos with the aim of identifying behavioral responses to aww noxious stimulus. For this purpose, a noxious mechanical stimulus and a control stimulus were applied. The recorded movements of the embryos were evaluated using the markerless pose estimation software DeepLabCut and manual observations. After the application of the noxious stimulus, a significant increase in beak movement was identified in 15- to 18-day-old embryos. In younger embryos, no behavioral changes related to the noxious stimulus were observed. The results indicate that noxious stimuli at the beak base evoke a nocifensive reaction in chicken embryos starting at embryonic day 15.

## Introduction

The behavior of birds can profoundly differ from the behavior of mammals, especially in terms of indications of pain^1^. For a long time, birds were not believed to feel pain^1^. At present, it is generally accepted that birds are capable of nociception and can feel pain^1,2^. Several studies have established that birds have mechanothermal, mechanical and thermal nociceptors with high stimulus thresholds^2,3^. Furthermore, peripheral and central processing of a potentially noxious stimulus in birds occurs in a similar manner to that in mammals^4^. Raja et al. define pain as an aversive experience of an individual that includes both sensory perception and emotional aspects^5^. This experience may be caused by a potential or actual lesion of the tissue^5^. Nociception, on the other hand, is described as the detection of a potentially damaging stimulus by primary sensory neurons and its processing in the nervous system^5,6^. The inability to communicate does not exclude the possibility that pain is felt, for example, by animals or neonates^1,5^. Another definition of pain more suitable for assessing pain in animals includes changes in species-specific behavior as a possible consequence of a painful experience^7^. Because pain is a subjective experience, its assessment is difficult in humans and is even more challenging in animals^1,5^. Detection and quantification of pain in animals involves inference from parameters associated with pain in humans^1^.

Birds show only subtle behaviors of discomfort or pain due to the disadvantage of showing weakness in a social group or as a prey species in general as well as the potential predominance of the flight reflex^8^. In addition, bird behavior varies greatly among species and individuals, making it necessary to closely examine the typical behavior of the observed individual. This makes it possible to assess deviations in typical behavior as a sign of pain^9^. Although pain-associated behavior is difficult to identify, its major advantage is that it can be observed immediately and noninvasively^3, 9^. This makes behavioral observation an essential part of a comprehensive pain assessment in birds.

Behavioral studies have been conducted in a variety of avian species^10^. Many of these studies used chickens (*Gallus gallus domesticus*) and evaluated nociceptive responses to procedures that are assumed to be painful or elicit discomfort^10,11^. The typical behavior of chicken embryos has long attracted scientific interest^10,11^. In the 1960s, the motility of chicken embryos was intensively studied. Movements and motility patterns, along with other aspects, were observed from days 3.5 to 20 of incubation^12-15^. In contrast, little is known about nociception in the chicken embryo or about nocifensive behavioral responses. According to current understanding, nociception in chicken embryos does not occur before the seventh day of incubation^16-18^.

The results presented are part of a comprehensive study investigating the developmental day at which chicken embryos are capable of nociception and pain perception. The aim of the present part III of the study was to evaluate the acute behavioral responses of chicken embryos at different developmental stages to a noxious mechanical stimulus. The markerless pose estimation software DeepLabCut (DLC) and manual observations were used to analyze embryonic behavior^19-21^. In addition, cardiovascular^22^ and electrophysiological^23^ parameters were investigated in parts I and II of the comprehensive study.

## Results

### Beak movements in response to a noxious stimulus

To analyze the movements of chicken embryos, the markerless pose estimation software DLC was used. The angle (*Beak Angle*) and distance (*Beak Distance*) between the upper and lower beak were calculated to reflect the opening of the beak as a potential response to a noxious mechanical stimulus applied at the base of the beak. The mechanical stimulation of the beak led to a change in the beak position at embryonic day (ED) 9 and ED12; thus, evaluation with DLC was distorted and could not be interpreted. At ED13 and ED14, *Beak Distance* did not differ between any time intervals during the two minutes after the control touch stimulus (hereafter, *Post Touch*) and the time intervals during the two minutes after the noxious pinch stimulus (hereafter, *Post Pinch*) (Supplementary Fig. 1). At ED15, significant increases in *Beak Distance* as a response to *Pinch* were detected (Fig. 1). Additionally, in ED15 embryos, beak movements *Post Pinch* increased significantly over the first 120 seconds compared to *Baseline Pinch* and over the first 90 seconds compared to *Post Touch*. On ED16, ED17 and ED18, a significant increase in *Beak Distance* was observed over all time intervals *Post Pinch* compared to *Baseline Pinch* and *Post Touch*. The greatest increase in *Beak Distance* occurred during the first 30 seconds of *Post Pinch*. The group of ED18 embryos that received an injection of the local anesthetic lidocaine (ED18 w/ Lido) did not exhibit reduced beak movements compared to same-age embryos that did not receive analgesia (Supplementary Fig. 2). *Beak Distance* was still significantly increased in ED18 w/ Lido in the first 30 seconds of *Post Pinch* (p<0.0001).

**Figure 1.**
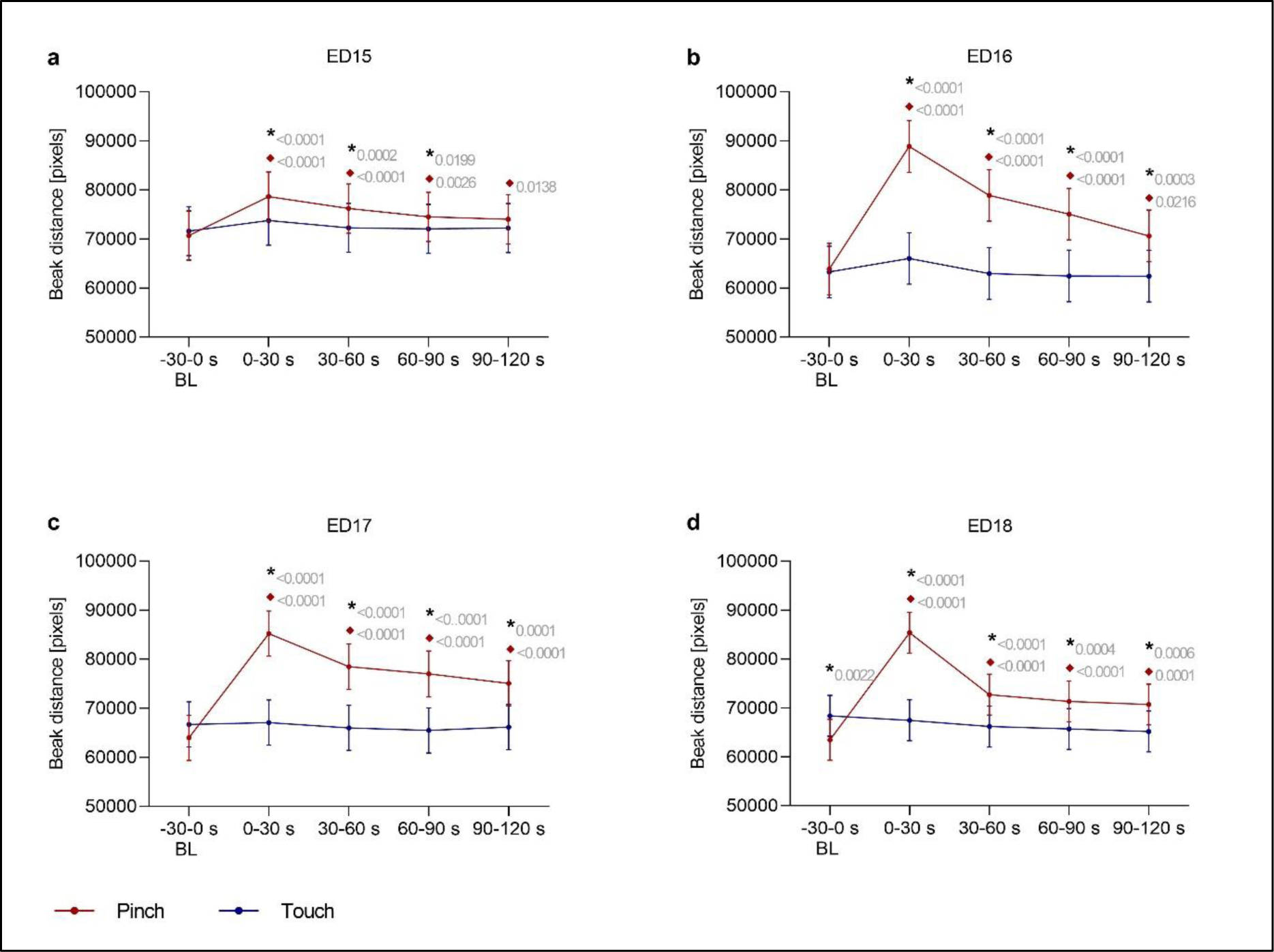
Beak Distance. This variable was defined as the distance between the upper and lower beak of embryos. It was measured at **a** ED15 (n=16), **b** ED16 (n=16), **c** ED17 (n=16) and **d** ED18 (n=15), before and after application of a control (*Touch*) or noxious stimulus (*Pinch*). The total distance in pixels across 30-second intervals (1500 frames) was evaluated. Plots show the estimated mean ± 95 % confidence intervals at the following 30-second intervals from Baseline (BL) to Post stimulation, with stimulation occurring at 0 s: -30–0, 0–30, 30–60, 60–90, and 90–120 seconds. Robust linear mixed effects were applied for all analysis. All contrasts (differences) between particular groups were assessed after model-fitting by the estimated marginal means with Tukey P value correction for multiple comparisons. *Touch*: blue; *Pinch*: red. * Significant difference between *Pinch* and *Touch*; ♦ Significant difference from baseline. P values shown.

*Beak Angle* results are displayed in the Supplementary Information (Supplementary Fig. 3). Briefly, *Beak Angle* showed a similar pattern of changes as *Beak Distance*. Additionally, significant increases in *Beak Angle* during *Post Pinch* were observed from ED15 onward.

### Head movements in response to a noxious stimulus

The medial eye corner was tracked to analyze the head movements of chicken embryos. Changes were particularly observed on ED13 and ED16 to ED18 in the first 30 seconds of *Post Pinch*. On these days, the embryos showed a significant increase in head movements after *Pinch* compared to after *Touch* (ED13: p=0.0254; ED16: p=0.0381; ED17: p=0.026; ED18: p<0.0001) and during *Baseline Pinch* (ED13: p=0.0256; ED16: p=0.0001; ED17: p<0.0001; ED18: p<0.0001). At ED12, head movements increased significantly at 30–60 seconds after *Pinch* compared to those 30–60 seconds after *Touch* (p=0.0372). At ED14, head movements also increased significantly in the first 30 seconds after the stimulus compared to those in the corresponding baseline period. These movements were observed after both stimuli (*Pinch*: p=0.0153; *Touch*: p=0.0069). In addition, a significant difference between head movements in response to *Pinch* and those in response to *Touch* was observed at 30— 60 seconds after the stimulus (p=0.0175). Head movements were significantly reduced in ED18 w/ Lido embryos in the first 30 seconds of *Post Pinch* compared to those of ED18 embryos in the same period (p<0.0001). Head movements on ED15 to ED18 are displayed in Fig. 2, while data on ED9, ED12 to ED14 and ED18 w/ Lido embryos is provided in the Supplementary Information (Supplementary Figs. 4 and 5).

**Figure 2.**
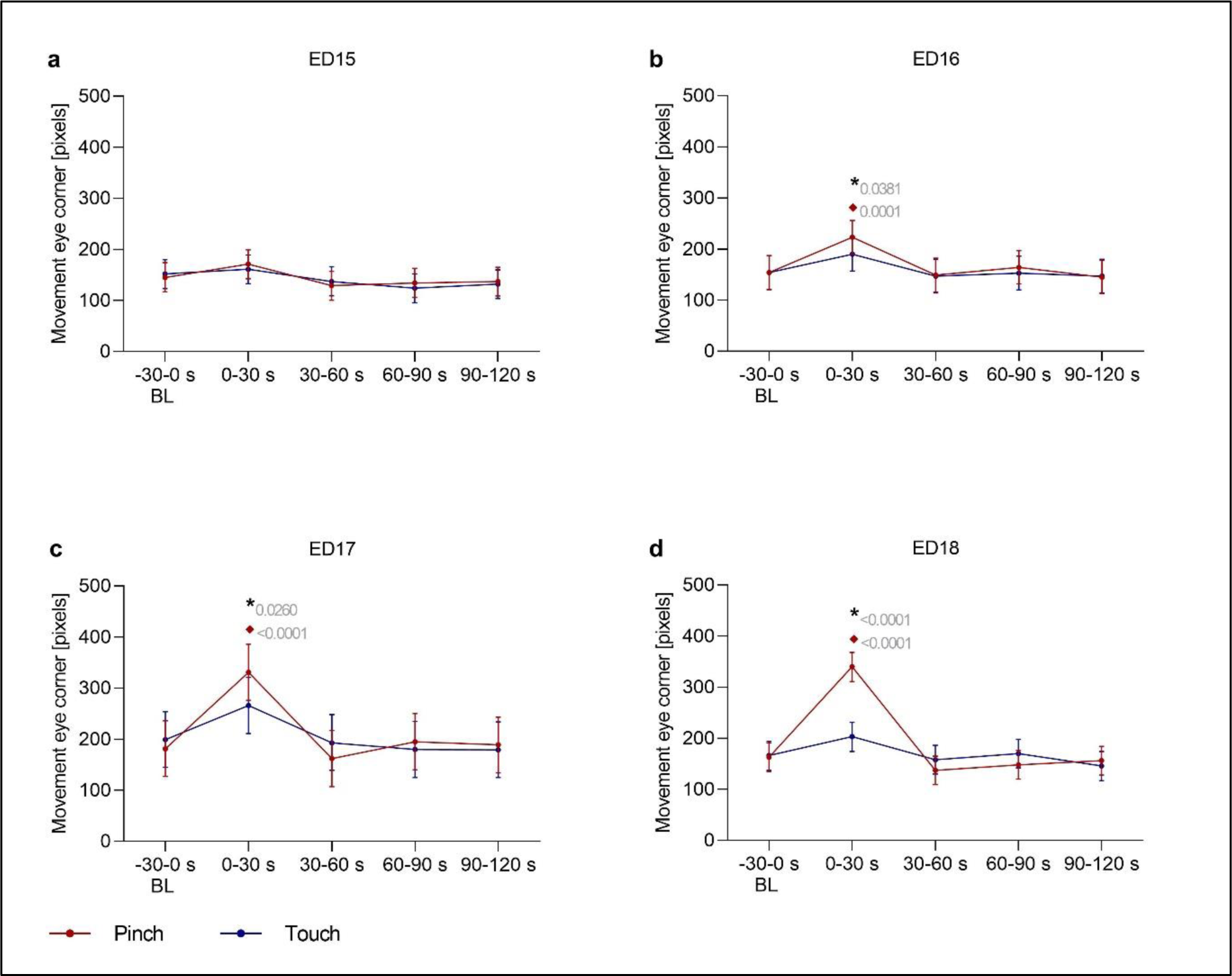
Eye Corner Movement. This variable was used to detect head movements of embryos at **a** ED15 (n=16), **b** ED16 (n=16), **c** ED17 (n=16) and **d** ED18 (n=15), before and after application of two stimuli (*Touch* and *Pinch*). The total distance in pixels across 30-second intervals (1500 frames) was evaluated. Plots show the estimated mean ± 95 % confidence intervals at the following 30-second intervals from Baseline (BL) to Post stimulation, with stimulation occurring at 0 s: -30–0, 0–30, 30–60, 60–90, and 90–120 seconds. Robust linear mixed effects were applied for all analysis. All contrasts (differences) between particular groups were assessed after model-fitting by the estimated marginal means with Tukey P value correction for multiple comparisons. *Touch*: blue; *Pinch*: red. * Significant difference between *Pinch* and *Touch*; ♦ Significant difference from baseline. P values shown.

**Figure 3.**
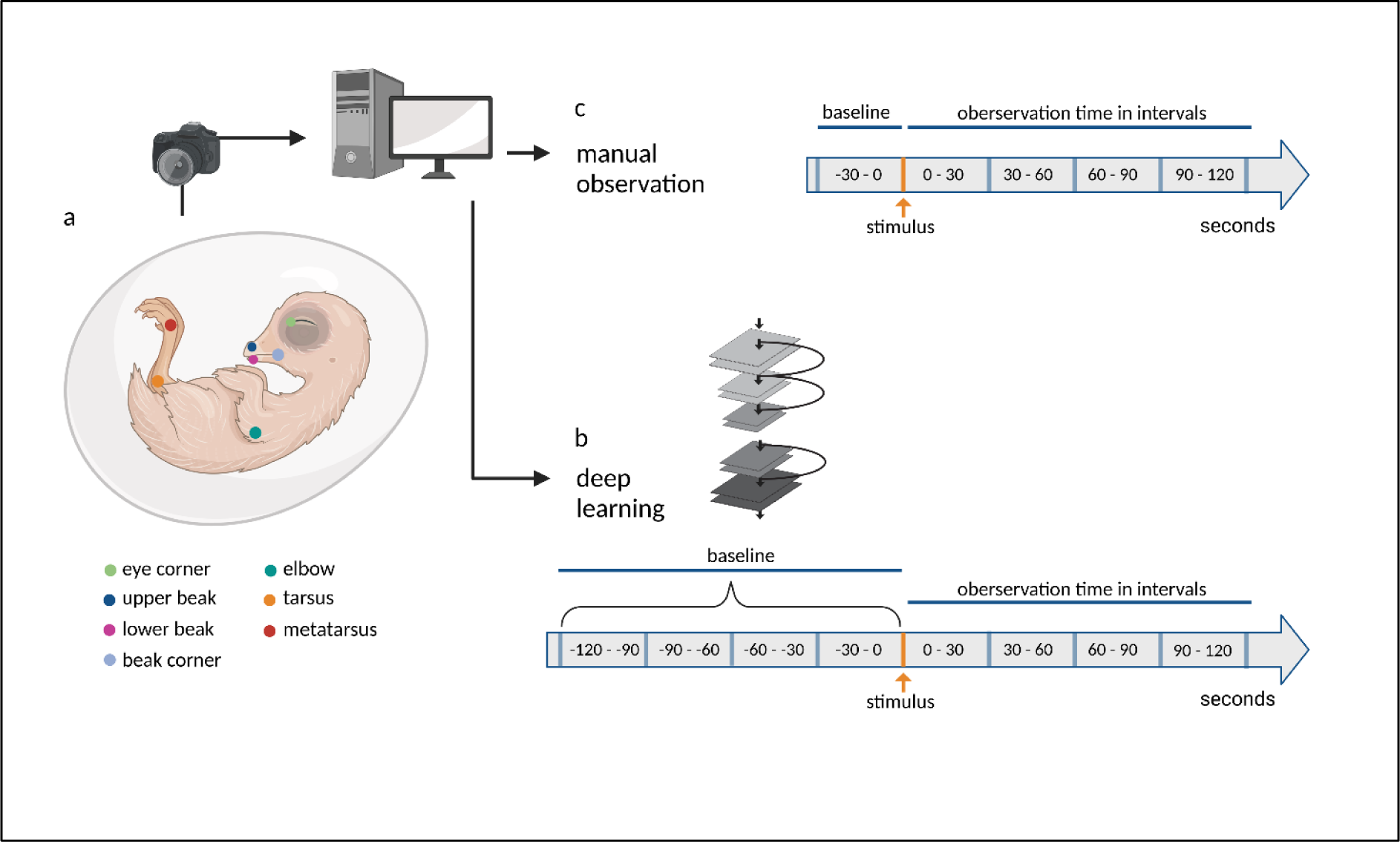
Flowchart of experimental procedures. **a** Recordings of the embryo were collected *in ovo,* and video data was transferred to a computer for editing. The body parts of chicken embryos tracked by DLC are labeled in the schema. **b** The neural network was trained and the video material was analyzed according to the timeline. **c** The video material was manual analyzed according to the timeline. (Created with BioRender.com).

### Limb movements in response to a noxious stimulus

To track limb movements, the movements of the *Elbow, Metatarsus* and *Tarsus* (ED9) were analyzed. Significant differences in limb movements between *Baseline Pinch* and *Post Pinch* and between *Post Pinch* and *Post Touch* were observed only on ED18 (Supplementary Figs. 6 and 7). An increase in elbow movements was observed between *Baseline Pinch* and *Post Pinch* (p=0.0023) as well as between *Post Pinch* and *Post Touch* (p=0.0096) during the first 30 seconds after the stimulus. Regarding the metatarsus movements, ED18 embryos showed a significant increase between *Baseline Pinch* and *Post Pinch* (p<0.0001) as well as between *Post Pinch* and *Post Touch* (p=0.0002) during the first 30 seconds after the stimulus. For ED18 w/ Lido embryos, no significant differences in limb movements were observed between *Baseline* and the first 30 seconds of *Post Stimulus*. There was also no significant difference between the ED18 embryos and the ED18 w/ Lido embryos. Other significant changes in limb movements were observed at specific time intervals over development.

### Characterization of beak movements in response to a noxious stimulus

In particular, DLC analysis identified changes in beak movement during *Post Pinch* in embryos from ED15 to ED18. To characterize beak movements in further detail, manual observations were performed. The focus of the manual observations was on four behaviors: *Beak Shift*, *Mandibulation*, *Beak Opening* and *Wide Beak Opening*. An overview of the percentage of animals that exhibited each behavior at specific time intervals is shown in Table 1. In addition, the counts of each behavior are shown in Supplementary Figs. 8–11.

**Table 1.** Percentage of chicken embryos showing beak movements. Overview of the percentage of chicken embryos that showed beak movements (*Beak Shift*, *Mandibulation*, *Beak Opening*, or *Wide Beak Opening*) during the 30 seconds before (*Baseline*) and 30 seconds after (*Post*) the stimulus.

| Amount of embryos [%] |  | ED9<br>n=10 |  | ED12<br>n=10 |  | ED13<br>n=10 |  | ED14<br>n=16 |  | ED15<br>n=16 |  | ED16<br>n=16 |  | ED17<br>n=16 |  | ED18<br>n=16 |  | ED18<br>w/ Lido<br>n=5 |  |
| --- | --- | --- | --- | --- | --- | --- | --- | --- | --- | --- | --- | --- | --- | --- | --- | --- | --- | --- | --- |
|  |  | *Touch* | *Pinch* | *Touch* | *Pinch* | *Touch* | *Pinch* | *Touch* | *Pinch* | *Touch* | *Pinch* | *Touch* | *Pinch* | *Touch* | *Pinch* | *Touch* | *Pinch* | *Touch* | *Pinch* |
| **Beak Shift** | *Baseline* | 0.0 | 0.0 | 0.0 | 0.0 | 10.0 | 30.0 | 18.8 | 25.0 | 31.3 | 0.0 | 18.8 | 25.0 | 25.0 | 31.3 | 25.0 | 6.3 | 40.0 | 40.0 |
|  | *Post* | 0.0 | 0.0 | 30.0 | 30.0 | 20.0 | 20.0 | 18.8 | 31.3 | 31.3 | 25.0 | 18.8 | 18.8 | 25.0 | 18.8 | 31.3 | 6.3 | 20.0 | 60.0 |
| **Mandibulation** | *Baseline* | 20.0 | 30.0 | 40.0 | 30.0 | 10.0 | 50.0 | 12.5 | 12.5 | 37.5 | 12.5 | 62.5 | 25.0 | 43.8 | 37.5 | 31.3 | 37.5 | 80.0 | 80.0 |
|  | *Post* | 30.0 | 20.0 | 50.0 | 60.0 | 50.0 | 40.0 | 56.3 | 56.3 | 62.5 | 81.3 | 68.8 | 93.8 | 62.5 | 87.5 | 68.8 | 87.5 | 80.0 | 60.0 |
| **Beak Opening** | *Baseline* | 10.0 | 0.0 | 10.0 | 10.0 | 10.0 | 10.0 | 6.3 | 0.0 | 0.0 | 6.3 | 0.0 | 6.3 | 6.3 | 6.3 | 0.0 | 0.0 | 0.0 | 20.0 |
|  | *Post* | 0.0 | 0.0 | 00.0 | 10.0 | 10.0 | 20.0 | 0.0 | 12.5 | 12.5 | 31.3 | 31.3 | 87.5 | 18.8 | 50.0 | 18.8 | 62.5 | 0.0 | 20.0 |
| **Wide Beak Opening** | *Baseline* | 0.0 | 0.0 | 0.0 | 0.0 | 10.0 | 0.0 | 0.0 | 6.3 | 0.0 | 0.0 | 0.0 | 0.0 | 0.0 | 0.0 | 0.0 | 0.0 | 20.0 | 0.0 |
|  | *Post* | 0.0 | 0.0 | 0.0 | 0.0 | 0.0 | 0.0 | 0.0 | 6.3 | 12.5 | 18.8 | 6.3 | 25.0 | 0.0 | 81.3 | 0.0 | 87.5 | 0.0 | 40.0 |

*Beak Opening* was rarely displayed during *Baseline* and was observed in only 10.0 % of animals from ED9 to ED18. *Beak Opening* was particularly rare on ED9 and ED12 to ED14. Before ED12, a maximum of 10.0 % of animals exhibited this behavior within a single time interval; up to ED14, a maximum of 20.0 % of animals exhibited this behavior within a single time interval. Starting from ED15, an increasing frequency (31.3 %) of *Beak Opening* was observed after the application of the noxious stimulus. At ED16, 87.5 % of embryos showed *Beak Opening* in the first 30 seconds of *Post Pinch*. Additionally, 50.0 % of ED17 embryos and 62.5 % of ED18 embryos showed this behavioral response to *Pinch*. During these days, at least twice as many embryos showed *Beak Opening* during *Post Pinch* as those during *Post Touch*.

*Wide Beak Opening*, characterized by visible tongue movement, was observed only sporadically during *Baseline* on all developmental days. This behavior was observed in only one animal each on ED13, ED14 and in ED18 w/ Lido embryos during baseline. Moreover, this specific beak movement was not observed during *Post Pinch* and *Post Touch* for ED9 to ED13 embryos and was observed only once during *Post Pinch* on ED14. On ED15 and ED16, this behavior was increasingly observed. A total of 18.8 % (ED15) and 25.0 % (ED16) of embryos exhibited *Wide Beak Opening* in the first 30 seconds of *Post Pinch*. A total of 81.3 % and 87.5 % of embryos on ED17 and ED18, respectively, showed more *Wide Beak Opening* in the first 30 seconds of *Post Pinch.* However, this behavior was never observed during *Post Touch* or corresponding baseline periods at these ages.

*Beak Shift* was observed from ED12 onward, but it did not appear to be associated with *Pinch*. *Mandibulation* was also observed across all embryonic days. Changes were observed in *Mandibulation* in all time in *Post Pinch and Post Touch* and regularly during both baseline periods.

Since *Beak Opening* and *Wide Beak Opening* were the most noticeable *Post Pinch* responses, the focus of comparisons with the additional control group that received local anesthetic (ED18 w/ Lido) was on these two movements, as the application of lidocaine reduced these behaviors. In the ED18 w/ Lido group, 40.0 % of the embryos reacted with *Wide Beak Opening* to the noxious mechanical stimulus; in the ED18 embryos without a lidocaine injection, 87.5 % exhibited this behavior. *Beak Opening* was observed in 20.0 % of the ED18 w/ Lido animals and 62.5 % of the untreated ED18 embryos. Neither *Mandibulation* or *Beak Shift* appeared to be associated with a specific reaction in any time interval, similar to embryos without lidocaine treatment. In other words, no noticeable increase or decrease in these behaviors was observed after a stimulus.

## Discussion

In this study, we investigated the movements of chicken embryos in response to a noxious stimulus at different developmental stages. We used DeepLabCut, a Python-based markerless pose estimation software, as well as manual observations to determine their responses.

Recently, the use of artificial intelligence and deep learning systems in behavioral studies has increased, and the availability of free software such as DLC allows such techniques to be used by researchers with less sophisticated programming experience^24-26^. In our study, we trained a model to provide satisfactory accuracy of tracking individual body parts on each embryonic day. One of the major advantages of using the markerless pose estimation software DLC is that it enables unbiased analysis. Calculations of distances are not based on subjective perception by an observer and are therefore quantifiable and reliable. Therefore, deep learning systems in general and DLC in particular offer a means of detecting and classifying behaviors that may not be detectible to the naked eye. However, the DLC analysis did not allow us to distinguish between types of beak movements. Thus, for better differentiation of beak movements, we added manual observation of these movements and identified four different patterns.

Pain behavior in general is influenced by a variety factors specific to the stimulus or the affected animal. For example, noxious agents can differ in duration (acute or chronic), source (somatic or visceral) and severity (mild to severe), each of which may provoke a different reaction^9,10,27^. Since behavioral responses vary extensively depending on the species and stimulus, any description is valid only for the specifically described case and cannot be transferred to another species without re-evaluation^10^. In our study, we applied an acute mechanical stimulus to the beak base of chicken embryos. The beak of chicken is known to be equipped with nocieptors^28^ and therefore represents a pain-sensitive area^11^. The beak has also been reported as the region in chicken embryos where the earliest response to stimuli is observed^29^. Chumak observed reflex movements in the form of flexions of the head on day 7 of incubation in response to pinpricks in the beak region, describing reflexes provoked by external stimuli (isolated movements of the head or wing) and spontaneous voluntary movements (involving generalized head, trunk, and limb movements)^29^.

Nociceptive reflexes have evolved as protective mechanisms^30^. A noxious stimulus is transmitted via peripheral nociceptors to the spinal cord and transmitted to motor neurons, resulting in muscle contraction and thus the nociceptive reflex^30-32^. Chumak reported more specific responses, including increased defensive movements, in chicken embryos at ED14/15, but characterized these responses as reflexive^29^. Hamburger and Oppenheim reported that coordinated movements appear around ED17^14^. Since our study was based solely on observations of movements by chicken embryos, a conclusion regarding whether the observed movements are reflexes or coordinated movements can therefore not be drawn.

We analyzed the movements of chicken embryos in response to a noxious stimulus applied to the beak from ED9 to ED18. Consistent with the assumption that a response to a stimulus is expected at the site of stimulus application, as was shown for well-innervated regions such as the beak^10^, our DLC data for *Beak Angle* and *Beak Distance* showed the most noticeable changes after the stimulus. Both parameters, *Beak Angle* and *Beak Distance,* quantified beak movements. A significant increase in beak movements was detected immediately after *Pinch* from ED15 to ED18. As the increase in beak movements during *Post Pinch* was significant compared to those during *Baseline Pinch* and *Post Touch*, we assumed that the increase in beak movements was a reaction to the noxious stimulus and was not a random movement of the chicken embryos.

Further differentiation of the movements through manual observation revealed that *Beak Opening* (starting on ED16) and *Wide Beak Opening* (starting on ED17) were recurring movements in response to the noxious stimulus. Individual, slow beak openings have been described in connection with the penetration of the air sac membrane shortly before hatching, at the end of day 18^14^. This description, however, does not match the rapid and clustered movements that we observed following the stimulus. Since these beak openings do not appear to be part of the typical behavior of chick embryos and markedly occurred only after a noxious stimulus, they may represent a nocifensive response by the embryo. Whether this can be interpreted as the presence of pain sensation remains unclear because an experience of pain presupposes consciousness^33^, and no indications can be made about this in the context of this part of the study.

Hamburger and Oppenheim also described a behavior that they called beak clapping, which involves rapid opening and closing of the beak in sequences that occurred at irregular intervals^14^. The description and random occurrence of this behavior matches *Mandibulation* in our study. Likewise, the movement was randomly observed across time intervals and had no clear connection to any of the stimuli. However, a similar behavior was observed in adult chickens as a response to low atmospheric pressure stunning before slaughter^34^. In this case, the mandibulation was discussed as a possible sign of reduced welfare or a physiological reaction to hypoxia^34^. As in the other studies, the embryos in our study underwent stress from the opening of the egg, the preparation, and the stimuli. Therefore, it is possible that *Mandibulation* is also a sign of stress in chicken embryos.

Application of the local anesthetic lidocaine did not yield a significant reduction in the beak movements of chicken embryos on ED18 according to the DLC analysis. However, in the manual observations, application of lidocaine reduced the percentage of embryos that responded to stimuli with *Wide Beak Opening* and *Beak Opening* by about half. Furthermore, local anesthetics are known to be effective in birds^35-37^ and can be used in chickens, e.g., for spinal anesthesia^38^ or a brachial plexus blockade^39^. However, there are no reliable empirical data regarding the mode of action of local anesthetics in chicken embryos. Additionally, we emphasize that only a small number of embryos were examined; thus, the results must be interpreted with caution. The inability of local anesthesia to reduce beak movements could also stem from the injection of lidocaine, which itself constitutes a noxious stimulus. In addition, numbness in the beak due to local anesthesia could have led to behavioral changes^40^. This is supported by the fact that head movements were significantly reduced by applying lidocaine at ED18.

Overall, stress could not be completely eliminated within the experimental setup; thus, its potential influence on behavior must be considered. The fenestrated egg does not represent a completely typical environment for the embryo because of the increased exposure to environmental influences, such as light. Additionally, the invasiveness of the preparation itself can induce stress, which is known to alter the behavior of birds^9^. We attempted to reduce external influences by standardizing the temperature and humidity during the experiments and adjusting them to match the typical incubation conditions as closely as possible. However, since direct access to the embryo was necessary for stimulation, and the embryo had to be visible to assess responses, some stressors were unavoidable.

We were also interested in whether limb movements changed after the noxious stimulus; however, we did not detect any overarching pattern until ED17. Occasional significant differences in limb movements during *Post Pinch* compared to those during *Baseline Pinch* or *Post Touch* were inconsistent over several EDs or time intervals and are therefore likely due to random movements, which have been described previously in^12-14,41-45^. Hamburger and Oppenheim stated that before ED15, the observed leg motility was not connected to any sensory input but appeared randomly due to autonomous cell discharges^15^. Wu et al. counted unilateral and bilateral simultaneous limb movements and found one maximum of movements between ED10 and ED13 for the former and two maxima on ED13 and ED17 for the latter^46^. In the present study, we detected a significant increase in elbow and metatarsal movements during the first 30 seconds of *Post Pinch* compared to those during the first 30 seconds of *Baseline Pinch* and *Post Touch* on only ED18, suggesting that these movements may represent an actual response to the noxious stimulus.

## Conclusion

We observed the movements of chicken embryos from ED9 to ED18 before and after noxious stimulation. During *Post Pinch*, the observed movement changes in ED15 to ED18 embryos were most likely a response to the noxious mechanical stimulus and can therefore be interpreted as nocifensive behavior. The results of our current movement analysis in combination with the corresponding results of the cardiovascular changes^22^ and the evaluation of the onset of physiological neuronal signals^23^ in chicken embryos during this developmental period provide valuable information that enhances our understanding of the development of nociception and pain perception in chicken.

## Material and Methods

### Animals and incubation

Chicken embryos from ED9 to ED18 were analyzed. An overview of the experimental groups is provided in Table 2. Fertilized Lohman Selected Leghorn eggs were obtained from the Technical University of Munich (TUM) Animal Research Centre, Thalhausen. Eggs were disinfected (Röhnfried Desinfektion Pro, Dr. Hesse Tierpharma GmbH & Co. KG, Hohenlockstedt Germany), weighed and stored in a refrigerator at 15 °C until use. The maximum storage time from the day of laying until the start of the incubation was seven days. Before incubation, the eggs were placed at room temperature for 24 hours. On the day of incubation, eggs were transferred at 8:30 am into a standard incubator (HEKA Favorit-Olymp 192 Spezial, HEKA-Brutgeräte, Rietberg, Germany) and incubated under the following conditions: 37.8 °C temperature and 55 % humidity. The eggs were turned six times a day until fenestration on ED3. The first day of incubation was defined as ED0.

**Table 2.** Number of chicken embryos. Overview of the number of chicken embryos analyzed on each embryonic day and the sex distribution.

|  | ED9 | ED12 | ED13 | ED14 | ED15 | ED16 | ED17 | ED18 | ED18<br>w/ Lido |
| --- | --- | --- | --- | --- | --- | --- | --- | --- | --- |
| Amount of embryos (n) | 10 | 10 | 10 | 16 | 16 | 16 | 16 | 16 | 5 |
| Sex<br>male/female | 5/5 | 3/7 | 5/5 | 9/7 | 7/9 | 7/8 | 7/9 | 7/9 | 2/3 |

On ED3, eggs were placed horizontally for two minutes, and 5–7 ml of albumin was withdrawn through a small hole at the pointed pole using a cannula. A small window was cut in the top of the eggshell, and 0.5 ml of penicillin‒streptomycin (10 000 units penicillin, 10 mg streptomycin/ml, P4333 – 100 ml, Sigma‒Aldrich, St. Louis, USA) were added. Eggs were sealed with plastic film and tape. With the eggs in a horizontal position, the incubation proceeded until the desired embryonic day^47^.

At the end of the experiments, the embryos were euthanized by an intravenous injection of pentobarbital-sodium (Narcoren, 16 g/100 ml, Boehringer Ingelheim Vetmedica GmbH, Ingelheim am Rhein, Germany; ED9: 0.05 ml, ED12 to ED15: 0.1 ml, and ED16 to ED18: 0.2 ml), followed by decapitation. Afterwards, the sex of ED12 to ED18 embryos was identified macroscopically by the determination of the gonads. For ED9 embryos, sexing was performed with PCR of genomic DNA samples isolated from pectoral and wing muscle. Screening was performed according to an established protocol^48^ using primers targeting the Z chromosome [5’ AAGCATAGAAACAATGTGGGAC 3’ (forward) and 5’ AACTCTGTCTGGAAGGACTT 3’ (reverse)] and female-specific primers targeting the W chromosome [5’ CTATGCCTACCACMTTCCTATTTGC 3’ (forward) and 5’ AACTCTGTCTGGAAGGACTT 3’ (reverse)]. The expected lengths of the DNA fragments were 250 bp and 375 bp, respectively, for female embryos and 250 bp for male embryos. An overview of the sex ratio on each ED is shown in Table 2.

### Preparation process

All experiments were performed between 9:00 am and 7:30 pm by the same two persons to standardize the procedure. To keep the environmental conditions as similar as possible to typical brooding conditions, experiments were conducted in a special heated chamber. The chamber was equipped with a heat mat (ThermoLux Wärmeunterlage, Witte + Sutor GmbH, Murrhardt, Germany), heat lamp (Wärmestrahlgerät, Taschenlampenwerk ARTAS GmbH, Arnstadt, Germany) and an air humidifier (Series 2000 Luftbefeuchter HU4811/10R1, Philips, Amsterdam, Netherlands). Humidity was kept at a constant level at 55.5 % ± 4.5. Additionally, the eggs were embedded in warm (38.0 °C) Armor Beads (Lab Armor Beads^TM^, Sheldon Manufacturing, Cornelius, USA). In this manner, the inner egg temperature was kept at 37.9 °C ± 0.9 during the entire experiment. To observe the entire embryo, the window in the eggshell was enlarged. Next, the chorioallantoic membrane (CAM) was carefully cut open and removed from the field of view. If necessary, blood vessels were ligated to prevent bleeding. However, to the extent possible, ligating or cutting vessels was avoided to prevent disruption of blood circulation. To gain access to the embryo and improve visibility, the amnion was carefully opened. A Desmarres lid retractor (Fuhrmann GmbH, Much, Germany) was carefully placed underneath the beak of the embryo to ensure beak visibility. In the case of ED9 embryos, a small wire loop was used.

### Experimental setup

All experiments were filmed with a camera (Panasonic LUMIX DC-G110V with a Panasonic Lumix G 30 m lens, Matsushita Electric Industrial Co., Ltd., Osaka, Japan; for ED9 to ED16: HOYA SUPER PRO1 Revo Filter SMC Cir-PL, Kenko Tokina Co, Ltd., Tokyo, Japan) with a frame rate of 50 frames per second.

After preparation, a resting period of three minutes was allotted. Baseline behavior was recorded for two (ED15 to ED18) or three (ED9 to ED14) minutes; subsequently, two stimuli were applied in a randomized order. The stimuli used were a noxious mechanical stimulus (*Pinch*) using a manual instrument and a light touch (*Touch*) as a negative control. Both were applied at the base of the beak. For ED15 to ED18 embryos, a mosquito clamp (Fine Science Tools, Foster City, USA) was used to administer the stimulus. To better monitor the applied force, a mosquito clamp combined with an analgesia meter (Rodent Pincher Analgesia Meter, Bioseb, Vitrolles, France) was used for experiments conducted with ED12 to ED14 embryos. Stimulus 1 (*Pinch* or *Touch*) was administered and followed by an observation duration of three minutes. After a second baseline period, stimulus 2 (*Touch* or *Pinch*) was administered, followed by another three minutes of observation. Because of their small size, microsurgical anatomical forceps (Fine Science Tools, Foster City, USA) had to be used to administer the stimulus to ED9 embryos. An additional group of ED18 embryos (ED18 w/ Lido) was injected with 0.02 ml of lidocaine (Xylocitin^®^ 2 %, Mibe GmbH Arzneimittel, Brehna, Germany) in the upper and lower beak region five minutes before the first baseline. Experiments were then performed according to the above protocol.

### Analyses: Hardware, software and statistical analyses

All videos were edited in the same way using the “daVinci Resolve” software (Blackmagic Design Pty. Ltd, Port Melbourne, Australia) before analysis. For each embryo, four single videos were cut referring to the sections of the experimental design: *Baseline Pinch*, *Baseline Touch*, *Post Pinch* and *Post Touch*.

### DeepLabCut

To track body parts of the embryo, the markerless pose estimation software DLC (version 2.2.1.1)^19,21^ was used on a computer (MSI MAG Infinite 11TC-1222AT, Intl Core i7–11700F, 16 GB RAM, nVidia GeForce RTX3060). The neural network was trained for each ED individually with video footage according to the protocol provided by the developers^21^. Manual labeling was always performed by the same person. The training was performed with the default settings and using a ResNet-50-based neural network^49,50^. A test error below 8.5 was obtained for every ED. After the model training was completed, the four experimental videos (*Baseline Pinch, Baseline Touch, Post Pinch,* and *Post Touch*) were analyzed for each embryo. For each labeled body part, DLC created three outputs for each frame of the video: an x coordinate, a y coordinate, and a likelihood value. These values were analyzed with custom-written code using MATLAB (MATLAB Version: 9.12.0.1927505 (R2022a) Update 1, MathWorks). In all cases, a likelihood value cutoff of 0.75 was used.

### Visualization of the data clusters

In the analysis, the focus was on the following body parts:

– Beak

– Head

– Limbs

– Stationary points on the egg, the Desmarres lid retractor, and the wire loop (for ED9) were used as a reference control.

As a first step, the labeled data clusters for each analyzed body part were visualized in the x-y coordinate space. This enabled refinement of the dataset through identification of outliers or mislabeled body parts. The videos were then checked for errors, and if any real outlier was found in a frame, its value was manually excluded.

### Distance between the upper and lower beak

The distance between the upper and lower boundaries of the beak was calculated in terms of the Euclidian distance between two points:

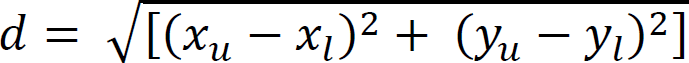

where *x*_*u*_ is the x coordinate of the upper beak label, *x*_*l*_ is the x coordinate of the lower beak label, *y*_*u*_ is the y coordinate of the upper beak label, and *y*_*l*_ is the y coordinate of the lower beak label. The Euclidian distance was calculated (in pixels) for every frame of the video.

### Angle between the upper and lower beak

The angle between the upper and lower beak was computed by calculating the angle between two lines *P*_0_ to *P*_1_ and *P*_0_ to *P*_2_, where *P*_0_ is the fulcrum between the beak parts, *P*_1_is the upper beak point and *P*_2_is the lower beak point. The angle was then calculated as follows:

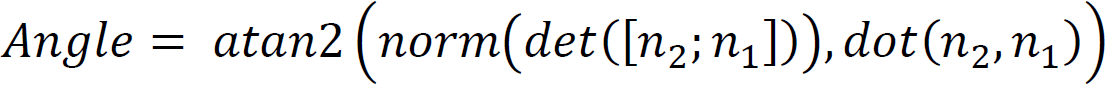

where *atan*2 is the four-quadrant inverse tangent, *det* is the matrix determinant, *dot* is the dot product, and *n*_2_, *n*_1_ are the Euclidean normalized vectors for *P*_0_ to *P*_1_and *P*_0_ to *P*_2_, respectively. The angle between the upper and lower beak was calculated for all frames of the video in radians and then converted to degrees.

### Movement

The movement of the body parts of interest was calculated in terms of the Euclidean distance between identical labels across consecutive frames:

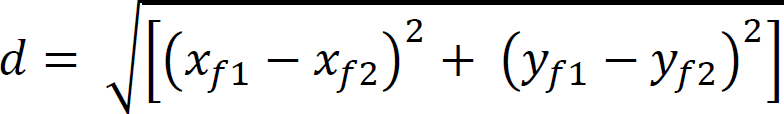

where *x*_*f*1_ is the x coordinate in frame 1, *x*_*f*2_ is the x coordinate in frame 2, *y*_*f*1_ is the y coordinate in frame 1, and *y*_*f*2_ is the y coordinate in frame 2. The distances were calculated for all consecutive frames. From ED12 to ED18, movements of the medial eye corner, elbow and metatarsus were analyzed. For the body movements on ED9, the tarsus (instead of the metatarsus) was used to assess leg movement, as the tissue of the metatarsus was translucent and prone to errors in tracking.

To simplify the analyses, 30-second intervals were evaluated. For each parameter, i.e., *Beak Distance*, *Beak Angle*, *Movement Eye Corner*, *Movement Elbow*, and *Movement Metatarsus*, the sum of the 1500 frame values of the interval was calculated. In Post Stimulus, this resulted in four intervals: 0–30, 30–60, 60–90, and 90–120 seconds. The beginning of the first poststimulus interval was defined as the moment from which the clamp was no longer in contact with the beak. The median of the four 30-second intervals prior to the stimulus was considered the baseline. Missing values, which arose after the exclusion of low likelihood values, were manually imputed. For each missing value series, the median was determined for half of the adjacent data and used in place of the missing value. If more than 5 % of the data in an interval were missing, the interval was excluded from the analysis. Due to a lack of visibility, one ED14 embryo and one ED18 embryo were completely excluded from the DLC analysis. A precise overview of the number of datasets ultimately included in the analysis is provided in Supplementary Table 1.

Due to the presence of repeated measures, generalized linear mixed effects models with the individual embryo as a random effect were chosen for analysis. Due to the violation of numerous model assumptions (normality of residual distribution, heteroscedasticity of residuals, heterogeneity of variances between groups and presence of outliers), only robust linear mixed-effects models were applied for all analyses (R package - robustlmm). All contrasts (differences) between particular groups were assessed after model-fitting by the estimated marginal means (R package - emmeans) with Tukey P value correction for multiple comparisons. The results with a P value < 0.05 were considered statistically significant. Data analysis was performed using R 4.2.1 (2022-06-23).

### Manual observation

The same video footage as used in the DLC analyses was used for manual observations. Since preliminary observations and data from the DLC analyses indicated that changes in beak position were frequent after *Pinch*, manual observations focused on beak movements. Four different patterns of beak movements were identified from the video material:

– *Beak Shift* – a small horizontal shift of the upper and lower beak against each other
– *Mandibulation* – a small vertical opening of the beak, often executed several times, and reminiscent of a chewing movement
– *Beak Opening* – single, swift, vertical opening of the beak
– *Wide Beak Opening* – single, wide, vertical opening of the beak; accompanied by a characteristic tongue movement

In an analogous approach to the one described above, the baseline and poststimulus observations were divided into intervals of 30 seconds. For manual observations, 30 seconds before the stimulus was used as a baseline. For each interval, the occurrences of the described beak movements were counted.

## Data availability

Raw data are available upon reasonable request to the corresponding author.

## Code availability

All MATLAB analysis code used in this study is available in a public GitHub repository: https://github.com/ondracej/dlcAnalysisEmbryo.

## Acknowledgments

This work was funded by the German Federal Ministry of Food and Agriculture (BMEL) based on a decision of the Parliament of the Federal Republic of Germany, granted by the Federal Office for Agriculture and Food (BLE; grant number 2821HS005).

The authors thank the scientific advisory board with Prof. Dr. Dr. Michael Erhard, Prof. Dr. Wolf Erhardt, Prof. Dr. Harald Luksch, Prof. Dr. Heidrun Potschka, Prof. Dr. Hans Straka and Dr. Britta Wirrer. In addition, the authors thank Marie-Louise Schmid, Dr. Johannes Fischer and Dr. Hicham Sid for their helpful support.

## Author contributions

Conceptualization: CB, JW, AMS, JR, TF, and BS. Data curation: SCS, JW, AMS, and JMO. Formal analysis: JMO and YZ. Funding acquisition: CB, TF, and BS. Investigation: SCS, LW, AMS, JW, and JR. Methodology: CB, JW, AMS, JR, SK, MA, TF, and BS. Project administration: CB. Resources: CB and BS. Supervision: CB, TF, and BS. Writing - original draft: SCS. Writing - review & editing: SCS, JW, AMS, SCS, JR, SK, MA, TF, BS, and CB. All authors have read and agreed to the published version of the manuscript.

