## Supplementary information for "Nociception in chicken embryos, Part III: Analysis of movements before and after application of a noxious stimulus"

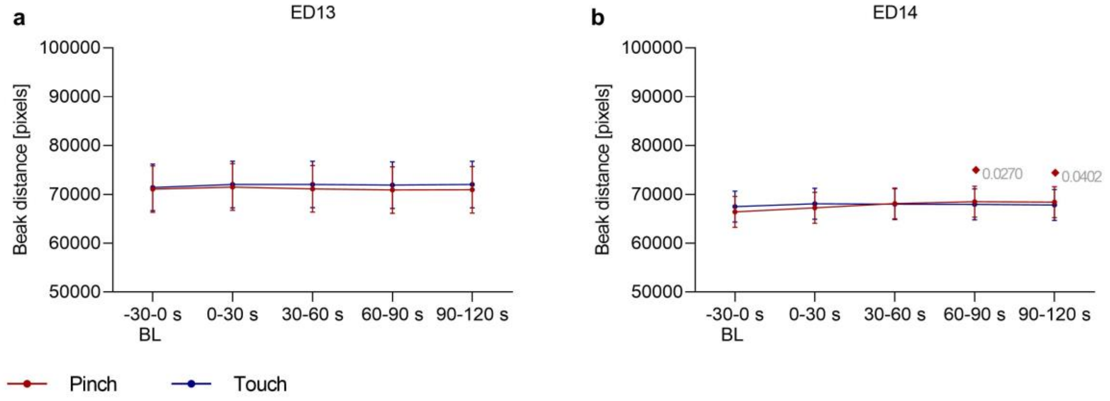


**Supplementary Fig. 1. Beak Distance.** This variable was defined as the distance between the upper and lower beak of embryos. It was measured at **a** ED13 (n=10) and **b** ED14 (n=15) before and after application of two stimuli (Touch and Pinch). The total distance in pixels across 30-second intervals (1500 frames) was evaluated. Plots show the estimated mean ± 95 % confidence intervals at the following 30-second intervals from Baseline (BL) to Post stimulation, with stimulation occurring at 0 s: -30–0, 0–30, 30–60, 60–90, and 90–120 seconds. Robust linear mixed effects were applied for all analysis. All contrasts (differences) between particular groups were assessed after model-fitting by the estimated marginal means with Tukey P value correction for multiple comparisons. *Touch*: blue; *Pinch*: red. ***** Significant difference between *Pinch* and *Touch*; ♦ Significant difference from Baseline. P values shown.

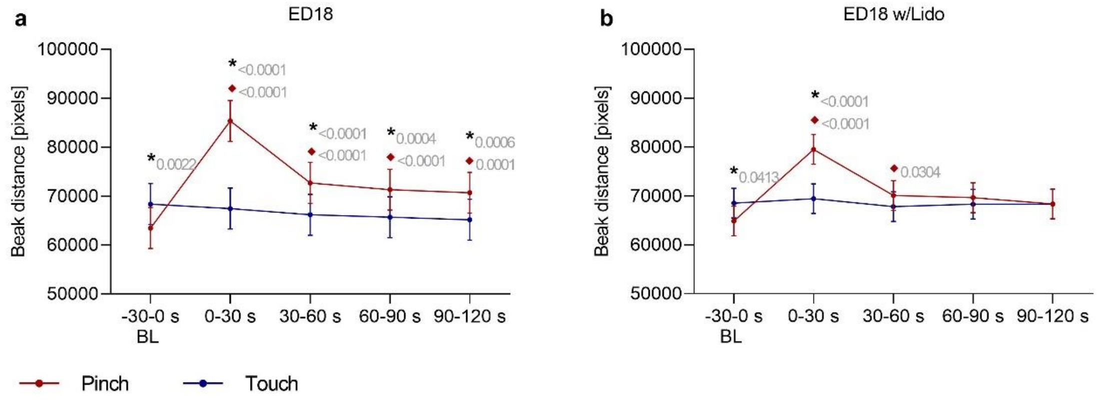


**Supplementary Fig. 2. Beak Distance Lidocaine** This variable was defined as the distance between the upper and lower beak of embryos. It was measured at **a** ED18 (n=15) and **b** ED18 w/Lido (n=5 before and after application of two stimuli (Touch and Pinch). The total distance in pixels across 30-second intervals (1500 frames) was evaluated. Plots show the estimated mean ± 95 % confidence intervals at the following 30-second intervals from Baseline (BL) to Post stimulation, with stimulation occurring at 0 s: -30–0, 0–30, 30–60, 60–90, and 90–120 seconds. Robust linear mixed effects were applied for all analysis. All contrasts (differences) between particular groups were assessed after model-fitting by the estimated marginal means with Tukey P value correction for multiple comparisons. *Touch*: blue; *Pinch*: red. ***** Significant difference between *Pinch* and *Touch*; ♦ Significant difference from baseline. P values shown.

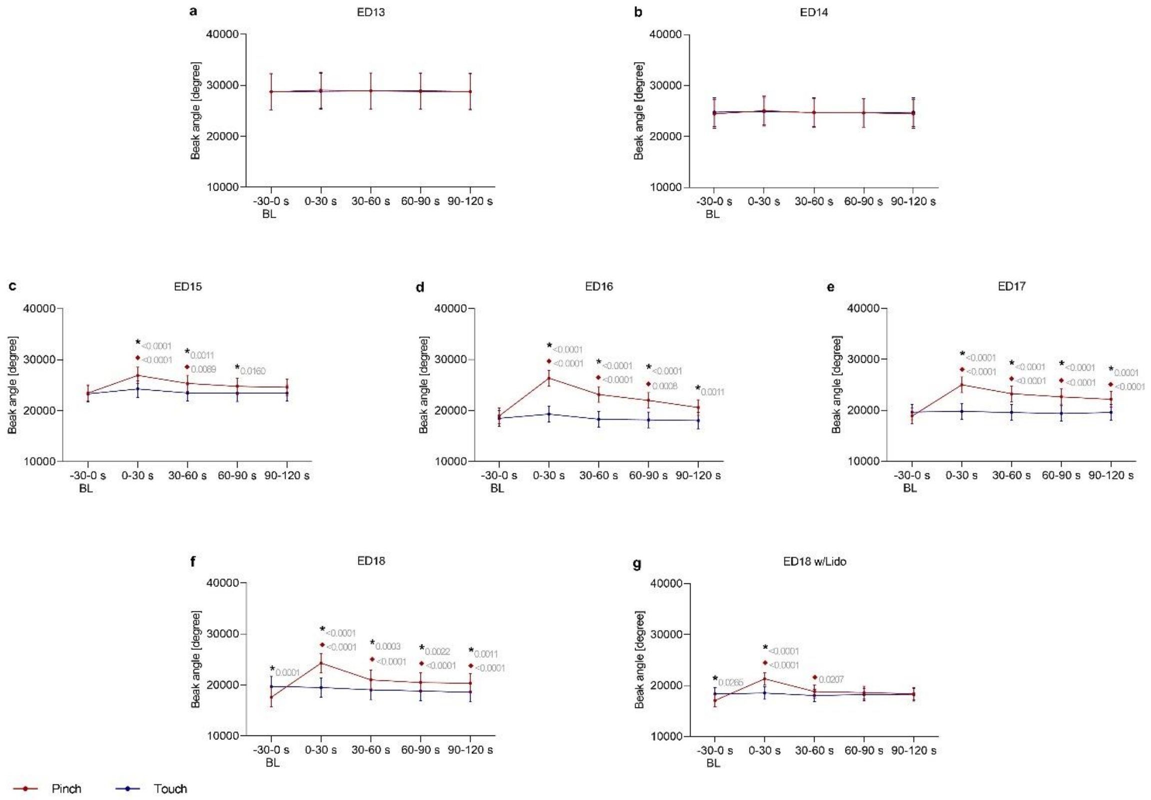


**Supplementary Fig. 3. Beak Angle.** This variable was defined as the angle between the beak corner, the upper and lower beak of embryos. It was measured at **a** ED13 (n=10), **b** ED14 (n=15), **c** ED15 (n=16), **d** ED16 (n=16,) **e** ED17 (n=16), **f** ED18 (n=16) and **g** ED18 w/Lido (n=5) before and after application of two stimuli (Touch and Pinch). The total distance in pixels across 30-second intervals (1500 frames) was evaluated. Plots show the estimated mean ± 95 % confidence intervals at the following 30-second intervals from Baseline (BL) to Post stimulation, with stimulation occurring at 0 s: -30–0, 0–30, 30–60, 60–90, and 90–120 seconds. Robust linear mixed effects were applied for all analysis. All contrasts (differences) between particular groups were assessed after model-fitting by the estimated marginal means with Tukey P value correction for multiple comparisons. *Touch*: blue; *Pinch*: red. ***** Significant difference between *Pinch* and *Touch*; ♦ Significant difference from baseline. P values shown.

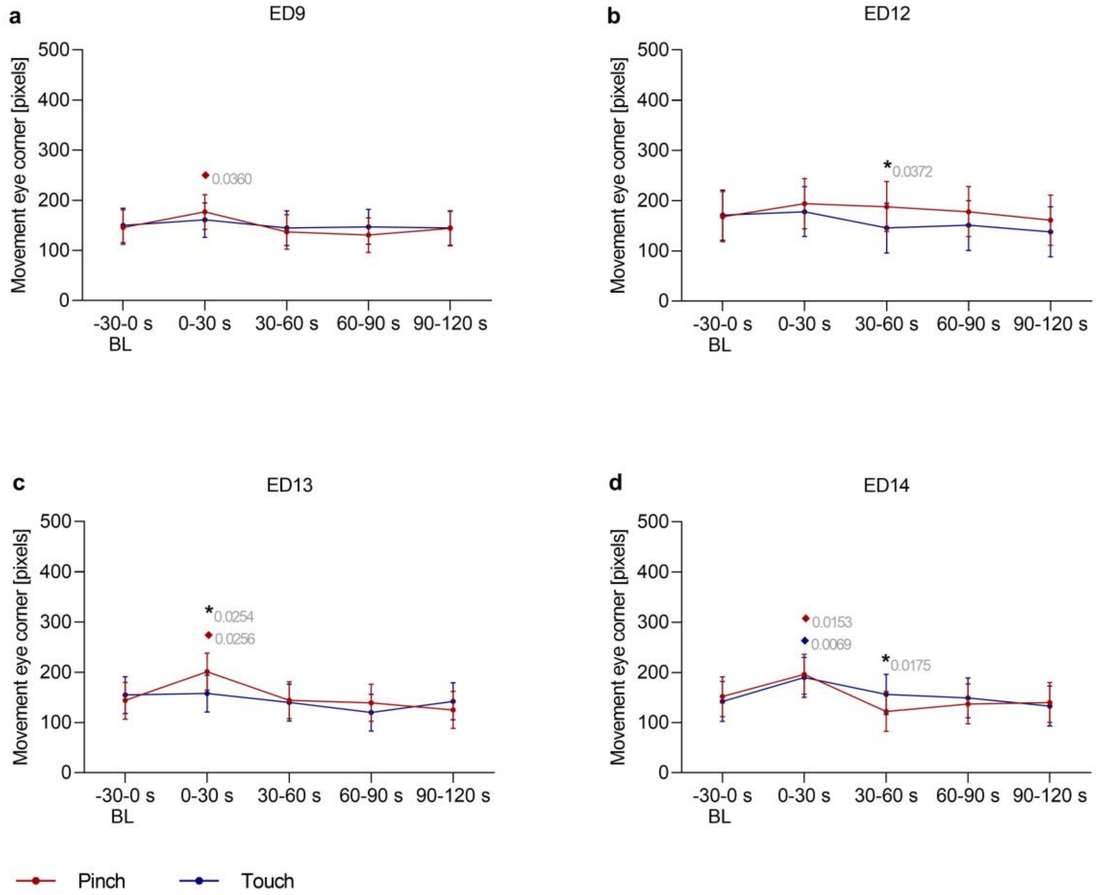


**Supplementary Fig. 4. Movement Eye corner.** This variable was used to detect head movements of embryos at **a** ED9 (n=10), **b** ED12 (n=10), **c** ED13 (n=10) and **d** ED14 (n=15) before and after application of two stimuli (Touch and Pinch). The total distance in pixels across 30-second intervals (1500 frames) was evaluated. Plots show the estimated mean ± 95 % confidence intervals at the following 30-second intervals from Baseline (BL) to Post stimulation, with stimulation occurring at 0 s: -30–0, 0–30, 30–60, 60–90, and 90–120 seconds. Robust linear mixed effects were applied for all analysis. All contrasts (differences) between particular groups were assessed after model-fitting by the estimated marginal means with Tukey P value correction for multiple comparisons. *Touch*: blue; *Pinch*: red. ***** Significant difference between *Pinch* and *Touch*; ♦ Significant difference from baseline. P values shown.

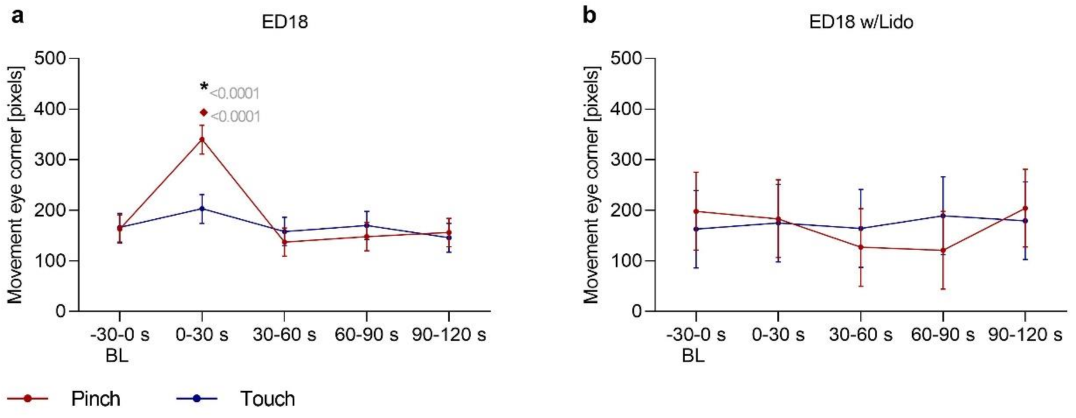


**Supplementary Fig. 5. Movement Eye corner Lidocaine** This variable was used to detect head movements of embryos at **a** ED18 (n=15) and **b** ED18 w/Lido (n=5) before and after application of two stimuli (Touch and Pinch). The total distance in pixels across 30-second intervals (1500 frames) was evaluated. Plots show the estimated mean ± 95 % confidence intervals at the following 30-second intervals from Baseline (BL) to Post stimulation, with stimulation occurring at 0 s: -30–0, 0–30, 30–60, 60–90, and 90–120 seconds. Robust linear mixed effects were applied for all analysis. All contrasts (differences) between particular groups were assessed after model-fitting by the estimated marginal means with Tukey P value correction for multiple comparisons. *Touch*: blue; *Pinch*: red. ***** Significant difference between *Pinch* and *Touch*; ♦ Significant difference from baseline. P values shown.

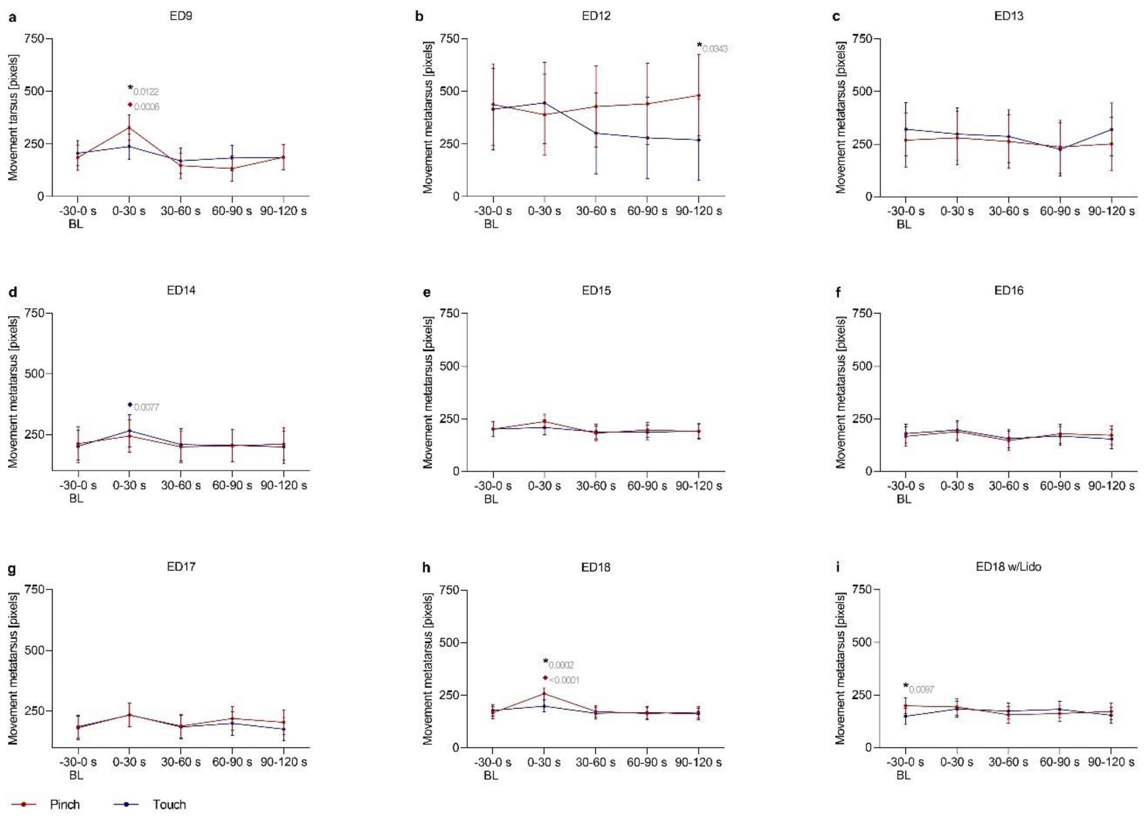


**Supplementary Fig. 6. Movement Metatarsus respectively Tarsus (ED9).** This variable was used to detect leg movements of embryos at **a** ED9 (n=10), **b** ED12 (n=10), **c** ED13 (n=10), **d** ED14 (n=15,) **e** ED15 (n=16), **f** ED16 (n=16), **g** ED17 (n=16), **h** ED18 (n=16) and **i** ED18 w/Lido (n=5) before and after application of two stimuli (Touch and Pinch). The total distance in pixels across 30-second intervals (1500 frames) was evaluated. Plots show the estimated mean ± 95 % confidence intervals at the following 30-second intervals from Baseline (BL) to Post stimulation, with stimulation occurring at 0 s: -30–0, 0–30, 30–60, 60–90, and 90–120 seconds. Robust linear mixed effects were applied for all analysis. All contrasts (differences) between particular groups were assessed after model-fitting by the estimated marginal means with Tukey P value correction for multiple comparisons. *Touch*: blue; *Pinch*: red. ***** Significant difference between *Pinch* and *Touch*; ♦ Significant difference from baseline. P values shown.

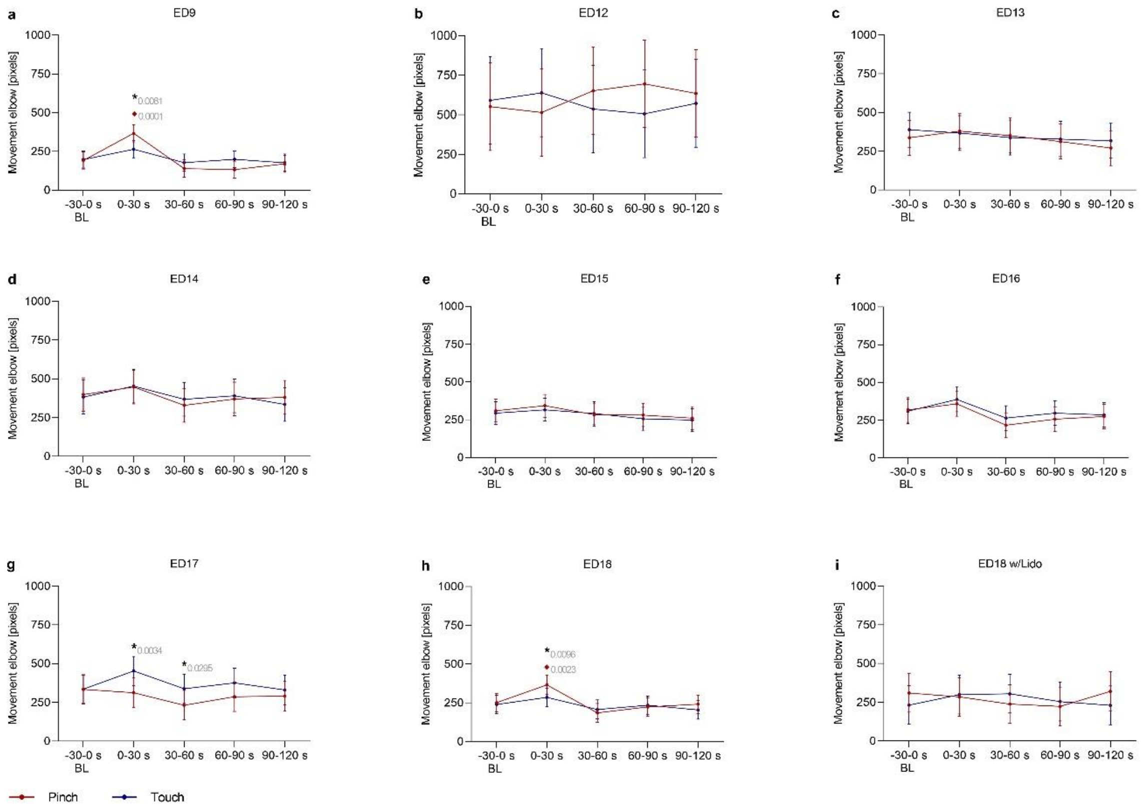


**Supplementary Fig. 7. Movement Elbow.** This variable was used to detect wing movements of embryos at **a** ED9 (n=10), **b** ED12 (n=10), **c** ED13 (n=10), **d** ED14 (n=15,) **e** ED15 (n=16), **f** ED16 (n=16), **g** ED17 (n=16), **h** ED18 (n=16) and **i** ED18 w/Lido (n=5) before and after application of two stimuli (Touch and Pinch). The total distance in pixels across 30-second intervals (1500 frames) was evaluated. Plots show the estimated mean ± 95 % confidence intervals at the following 30-second intervals from Baseline (BL) to Post stimulation, with stimulation occurring at 0 s: -30–0, 0–30, 30–60, 60–90, and 90–120 seconds. Robust linear mixed effects were applied for all analysis. All contrasts (differences) between particular groups were assessed after model-fitting by the estimated marginal means with Tukey P value correction for multiple comparisons. *Touch*: blue; *Pinch*: red. ***** Significant difference between *Pinch* and *Touch*; ♦ Significant difference from baseline. P values shown.

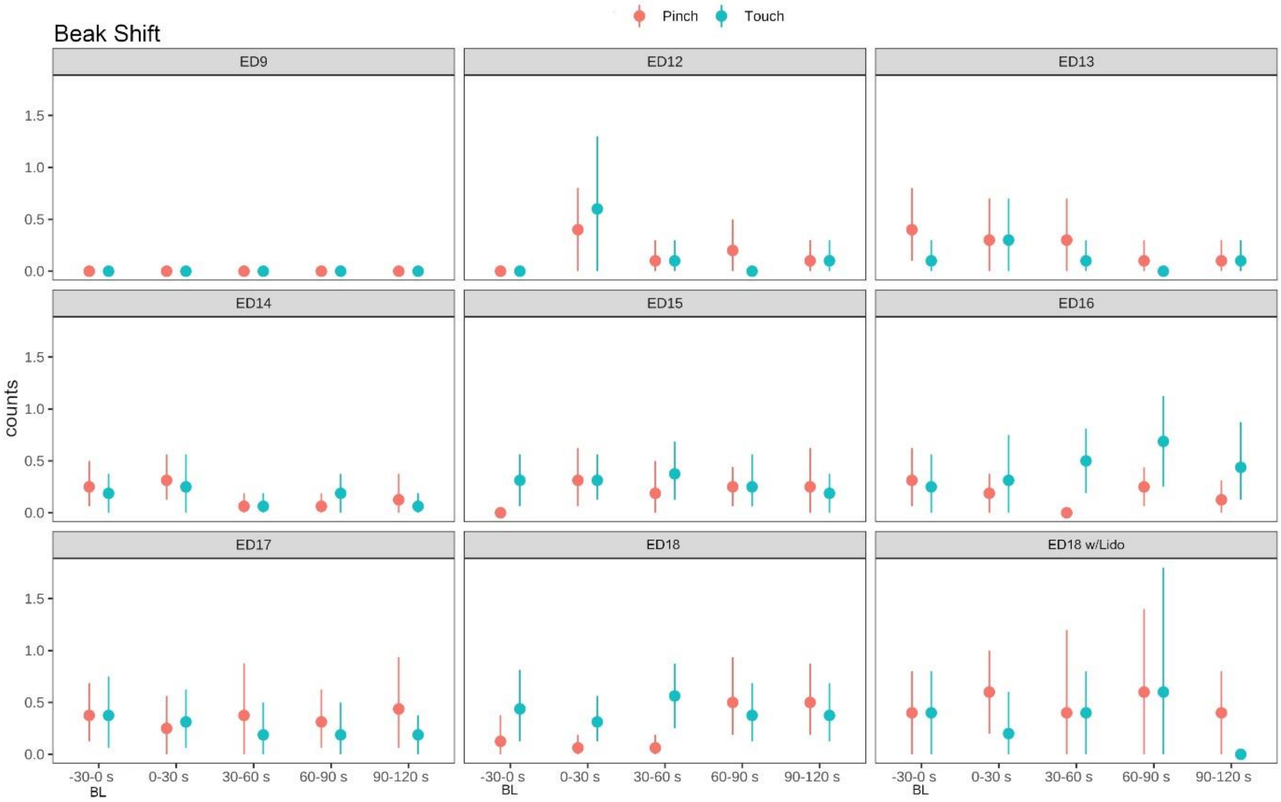


**Supplementary Fig. 8. Beak Shift.** The variable was used to detect the number of times (counts) embryos at ED9 (n=10), ED12 to ED13 (n=10), ED14 to ED18 (n=16) and ED18 w/ Lido (n=5) showed the behavior *Beak Shift* before and after stimulus (*Touch/Pinch*). A nonparametric bootstrap for obtaining confidence limits for the mean without assuming normality was used for the visualization of data.

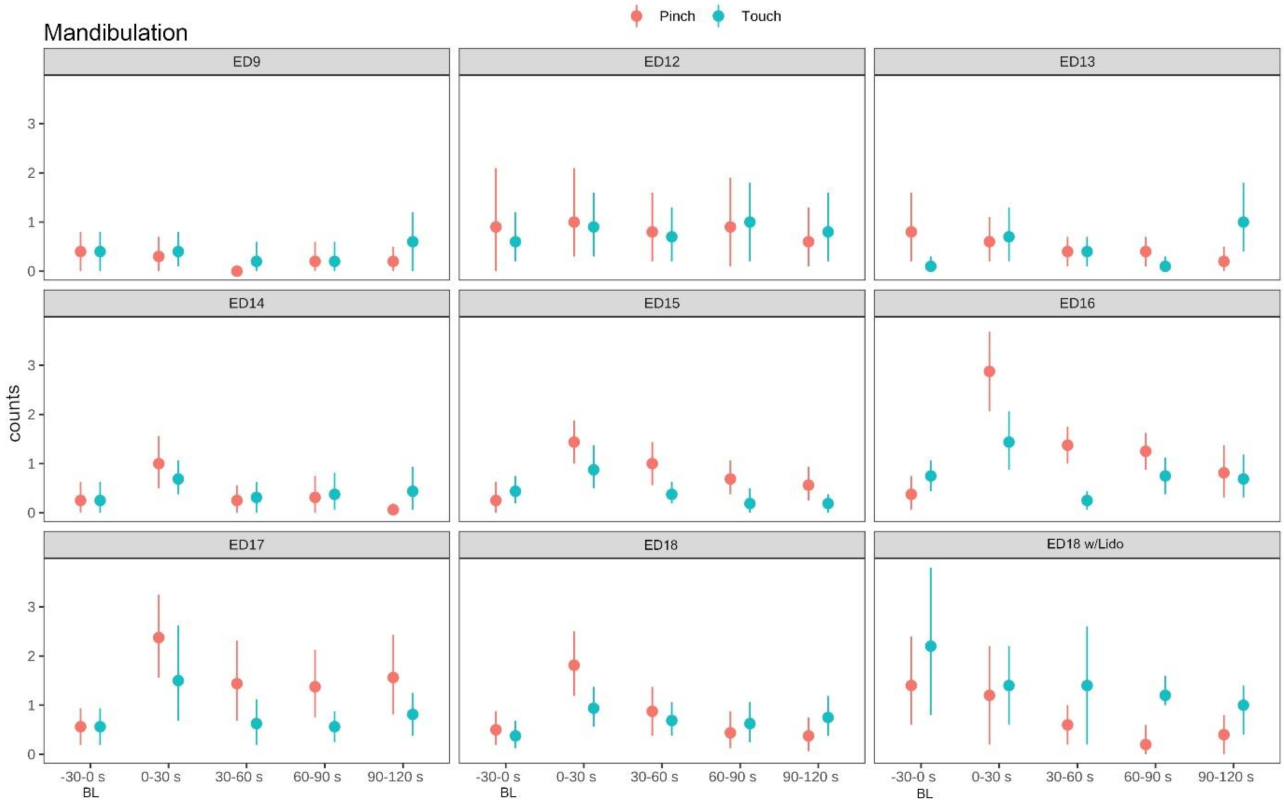


**Supplementary Fig. 9. Mandibulation.** The variable was used to detect the number of times (counts) embryos at ED9 (n=10), ED12 to ED13 (n=10), ED14 to ED18 (n=16) and ED18 w/ Lido (n=5) showed the behavior *Mandibulation* before and after stimulus (*Touch/Pinch*). A nonparametric bootstrap for obtaining confidence limits for the mean without assuming normality was used for the visualization of data.

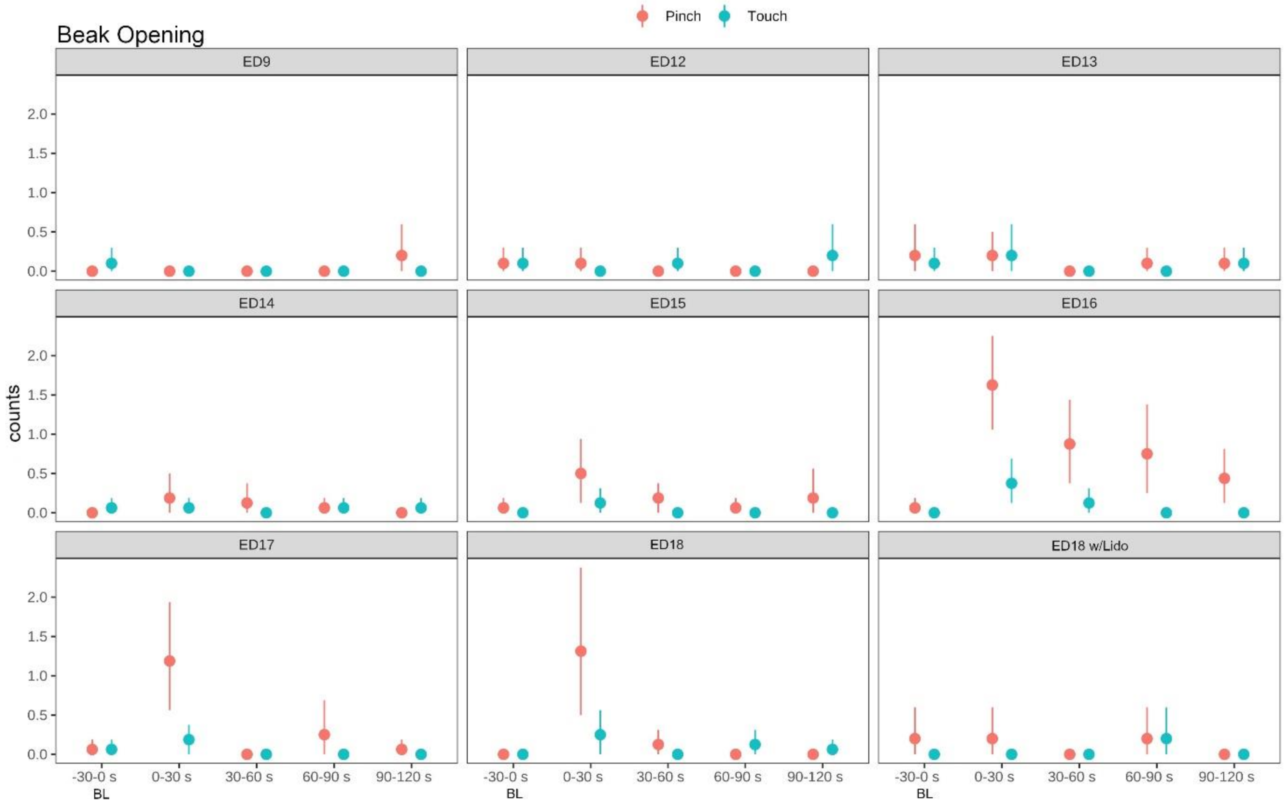


**Supplementary Fig. 10. Beak Opening.** The variable was used to detect the number of times (counts) embryos at ED9 (n=10), ED12 to ED13 (n=10), ED14 to ED18 (n=16) and ED18 w/ Lido (n=5) showed the behavior *Beak Opening* before and after stimulus (*Touch/Pinch*). A nonparametric bootstrap for obtaining confidence limits for the mean without assuming normality was used for the visualization of data.

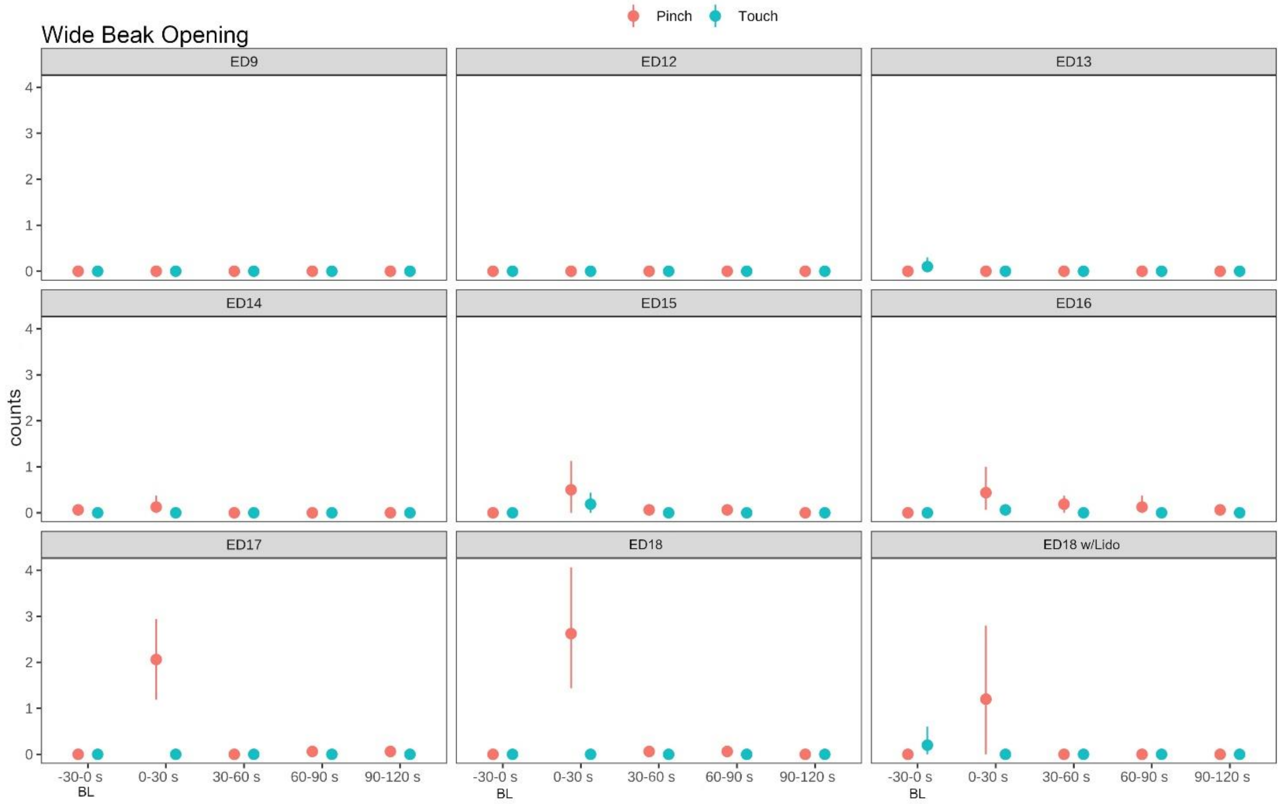


**Supplementary Fig. 11.** ***Wide Beak Opening.*** The variable was used to detect the number of times (counts) embryos at ED9 (n=10), ED12 to ED13 (n=10), ED14 to ED18 (n=16) and ED18 w/ Lido (n=5) showed the behavior *Wide Beak Opening* before and after stimulus (*Touch/Pinch*). A nonparametric bootstrap for obtaining confidence limits for the mean without assuming normality was used for the visualization of data.

**Supplementary Table 1**. Overview on the number of datasets from DLC included in the final analysis

| ED18 | | | | | | |
|---|---|---|---|---|---|---|
| | **Time Interval** | **Beak angle** | **Beak distance** | **Elbow** | **Eye corner** | **Metatarsus** |
| **Touch** | -30-0 sec | 15 | 15 | 15 | 15 | 14 |
| | 0-30 sec | 15 | 15 | 15 | 15 | 14 |
| | 30-60 sec | 15 | 15 | 15 | 15 | 14 |
| | 60-90 sec | 15 | 15 | 15 | 15 | 14 |
| | 90-120 sec | 15 | 15 | 15 | 15 | 14 |
| **Pinch** | -30-0 sec | 15 | 15 | 15 | 15 | 14 |
| | 0-30 sec | 15 | 15 | 15 | 15 | 14 |
| | 30-60 sec | 15 | 15 | 15 | 15 | 14 |
| | 60-90 sec | 15 | 15 | 15 | 15 | 14 |
| | 90-120 sec | 15 | 15 | 15 | 15 | 14 |
| **ED17** | | | | | | |
| | **Time Interval** | **Beak angle** | **Beak distance** | **Elbow** | **Eye corner** | **Metatarsus** |
| **Touch** | -30-0 sec | 16 | 16 | 16 | 16 | 16 |
| | 0-30 sec | 15 | 16 | 16 | 16 | 16 |
| | 30-60 sec | 16 | 16 | 15 | 16 | 16 |
| | 60-90 sec | 15 | 16 | 15 | 16 | 16 |
| | 90-120 sec | 15 | 16 | 15 | 16 | 16 |
| **Pinch** | -30-0 sec | 16 | 16 | 16 | 16 | 16 |
| | 0-30 sec | 15 | 16 | 15 | 16 | 15 |
| | 30-60 sec | 15 | 16 | 15 | 16 | 15 |
| | 60-90 sec | 16 | 15 | 15 | 16 | 15 |
| | 90-120 sec | 16 | 16 | 15 | 16 | 16 |
| **ED16** | | | | | | |
| | **Time Interval** | **Beak angle** | **Beak distance** | **Elbow** | **Eye corner** | **Metatarsus** |
| **Touch** | -30-0 sec | 16 | 16 | 16 | 16 | 16 |
| | 0-30 sec | 16 | 16 | 16 | 16 | 16 |
| | 30-60 sec | 16 | 16 | 16 | 16 | 16 |
| | 60-90 sec | 16 | 16 | 16 | 16 | 16 |
| | 90-120 sec | 15 | 16 | 16 | 16 | 16 |
| **Pinch** | -30-0 sec | 16 | 16 | 16 | 16 | 16 |
| | 0-30 sec | 16 | 16 | 16 | 16 | 16 |
| | 30-60 sec | 16 | 16 | 16 | 16 | 16 |
| | 60-90 sec | 16 | 16 | 16 | 16 | 16 |
| | 90-120 sec | 16 | 16 | 16 | 16 | 16 |

| ED15 | | | | | | |
|---|---|---|---|---|---|---|
| | Time Interval | Beak angle | Beak distance | Elbow | Eye corner | Metatarsus |
| Touch | -30-0 sec | 16 | 16 | 16 | 16 | 16 |
| | 0-30 sec | 16 | 16 | 16 | 16 | 16 |
| | 30-60 sec | 16 | 16 | 16 | 16 | 16 |
| | 60-90 sec | 16 | 16 | 16 | 16 | 16 |
| | 90-120 sec | 15 | 16 | 16 | 16 | 16 |
| Pinch | -30-0 sec | 16 | 16 | 16 | 16 | 16 |
| | 0-30 sec | 16 | 16 | 16 | 16 | 16 |
| | 30-60 sec | 16 | 16 | 16 | 16 | 16 |
| | 60-90 sec | 16 | 16 | 16 | 16 | 16 |
| | 90-120 sec | 16 | 16 | 16 | 16 | 16 |
| **ED14** | | | | | | |
| | **Time Interval** | **Beak angle** | **Beak distance** | **Elbow** | **Eye corner** | **Metatarsus** |
| **Touch** | -30-0 sec | 15 | 15 | 15 | 15 | 13 |
| | 0-30 sec | 15 | 15 | 15 | 15 | 13 |
| | 30-60 sec | 15 | 15 | 15 | 15 | 13 |
| | 60-90 sec | 15 | 15 | 15 | 15 | 13 |
| | 90-120 sec | 15 | 15 | 15 | 15 | 13 |
| **Pinch** | -30-0 sec | 15 | 15 | 15 | 15 | 13 |
| | 0-30 sec | 15 | 15 | 15 | 15 | 13 |
| | 30-60 sec | 15 | 15 | 15 | 15 | 13 |
| | 60-90 sec | 15 | 15 | 15 | 15 | 13 |
| | 90-120 sec | 15 | 15 | 15 | 15 | 13 |
| **ED13** | | | | | | |
| | **Time Interval** | **Beak angle** | **Beak distance** | **Elbow** | **Eye corner** | **Metatarsus** |
| **Touch** | -30-0 sec | 10 | 10 | 10 | 10 | 8 |
| | 0-30 sec | 10 | 10 | 10 | 10 | 9 |
| | 30-60 sec | 10 | 10 | 10 | 10 | 9 |
| | 60-90 sec | 10 | 10 | 10 | 10 | 9 |
| | 90-120 sec | 10 | 10 | 10 | 10 | 9 |
| **Pinch** | -30-0 sec | 10 | 10 | 10 | 10 | 9 |
| | 0-30 sec | 10 | 10 | 10 | 10 | 9 |
| | 30-60 sec | 10 | 10 | 10 | 10 | 9 |
| | 60-90 sec | 10 | 10 | 10 | 10 | 9 |
| | 90-120 sec | 10 | 10 | 10 | 10 | 9 |

| ED12 | | | | | | |
|---|---|---|---|---|---|---|
| | **Time Interval** | **Beak angle** | **Beak distance** | **Elbow** | **Eye corner** | **Metatarsus** |
| **Touch** | -30-0 sec | 10 | 10 | 9 | 10 | 8 |
| | 0-30 sec | 10 | 10 | 9 | 10 | 8 |
| | 30-60 sec | 10 | 10 | 9 | 10 | 8 |
| | 60-90 sec | 10 | 10 | 9 | 10 | 8 |
| | 90-120 sec | 10 | 10 | 9 | 10 | 8 |
| **Pinch** | -30-0 sec | 10 | 10 | 9 | 10 | 8 |
| | 0-30 sec | 10 | 10 | 9 | 10 | 8 |
| | 30-60 sec | 10 | 10 | 9 | 10 | 8 |
| | 60-90 sec | 10 | 10 | 9 | 10 | 8 |
| | 90-120 sec | 10 | 10 | 9 | 10 | 8 |
| **ED9** | | | | | | |
| | **Time Interval** | **Beak angle** | **Beak distance** | **Elbow** | **Eye corner** | **Tarsus** |
| **Touch** | -30-0 sec | 10 | 10 | 10 | 10 | 10 |
| | 0-30 sec | 10 | 10 | 10 | 10 | 10 |
| | 30-60 sec | 10 | 10 | 10 | 10 | 10 |
| | 60-90 sec | 10 | 10 | 10 | 10 | 10 |
| | 90-120 sec | 10 | 10 | 10 | 10 | 10 |
| **Pinch** | -30-0 sec | 10 | 10 | 10 | 10 | 10 |
| | 0-30 sec | 10 | 10 | 10 | 10 | 10 |
| | 30-60 sec | 10 | 10 | 10 | 10 | 10 |
| | 60-90 sec | 10 | 10 | 10 | 10 | 10 |
| | 90-120 sec | 10 | 10 | 10 | 10 | 10 |
| **ED18 w/ Lido** | | | | | | |
| | **Time Interval** | **Beak angle** | **Beak distance** | **Elbow** | **Eye corner** | **Metatarsus** |
| **Touch** | -30-0 sec | 5 | 5 | 5 | 5 | 5 |
| | 0-30 sec | 5 | 5 | 5 | 5 | 5 |
| | 30-60 sec | 5 | 5 | 5 | 5 | 5 |
| | 60-90 sec | 5 | 5 | 5 | 5 | 5 |
| | 90-120 sec | 5 | 5 | 5 | 5 | 5 |
| **Pinch** | -30-0 sec | 5 | 5 | 5 | 5 | 5 |
| | 0-30 sec | 5 | 5 | 5 | 5 | 5 |
| | 30-60 sec | 5 | 5 | 5 | 5 | 5 |
| | 60-90 sec | 5 | 5 | 5 | 5 | 5 |
| | 90-120 sec | 5 | 5 | 5 | 5 | 5 |
